# Helminth infection is associated with dampened cytokine responses to viral and bacterial stimulations in Tsimane hunter-horticulturalists

**DOI:** 10.1101/2021.09.29.462428

**Authors:** India Schneider-Crease, Aaron D. Blackwell, Thomas S. Kraft, Melissa Emery Thompson, Ivan Maldonado Suarez, Daniel K. Cummings, Jonathan Stieglitz, Noah Snyder-Mackler, Michael Gurven, Hillard Kaplan, Benjamin C. Trumble

**Affiliations:** Center for Evolution and Medicine, Arizona State University, Tempe, Arizona, USA; School of Life Sciences, Arizona State University, Tempe, Arizona, USA; Department of Anthropology, Washington State University, Pullman, Washington, USA; Department of Anthropology, University of California Santa Barbara, Santa Barbara, California, USA; Department of Anthropology, University of New Mexico, Albuquerque, New Mexico, USA; Tsimane Health and Life History Project, San Borja, Bolivia; Economic Science Institute, Chapman University, Orange, California, USA; Institute for Advanced Study in Toulouse, Toulouse, France; School of Human Evolution and Social Change, Arizona State University, Tempe, Arizona, USA

**Keywords:** Soil-transmitted helminths, viruses, bacteria, cytokine storms, eosinophilia, hypereosinophilia, immunomodulation

## Abstract

**Background:** Soil-transmitted helminth (STH) infections can catalyze immunological changes that affect the response to subsequent infections, particularly those that elicit strong inflammatory responses. As globalization heightens the risk that remote communities with high STH prevalence will encounter novel pathogens, understanding how STHs shape immune responses to these downstream infections becomes increasingly crucial.

**Methodology:** We worked with Tsimane forager-horticulturalists in the Bolivian Amazon, where STHs are prevalent. We tested whether STHs and eosinophil levels—most likely indicative of infection in this population—are associated with dampened immune responses to *in vitro* stimulation with H1N1 and lipopolysaccharide (LPS) antigens. Whole blood samples (n = 179) were treated with H1N1 vaccine and LPS and assayed for 13 cytokines (interferon gamma [INF-*γ*], interleukin [IL]-1*β*, IL-2, IL-4, IL-5, IL-6, IL-7, IL-8, IL-10, IL-12p70, IL-13, Granulocyte-macrophage colony-stimulating factor [GM-CSF], and Tumor necrosis factor-alpha [TNF-*α*]). We evaluated how STHs and eosinophil levels affected cytokine responses and T helper (Th) 1 and Th2-cytokine suite responses to stimulation.

**Results:** Infection with *Ascaris lumbricoides* was significantly (p ≤ 0.05) associated with lower response of some cytokines to H1N1 and LPS in women. Eosinophils were significantly negatively associated with some cytokine responses to H1N1 and LPS, with the strongest effects in women, and associated with a reduced Th1- and Th2-cytokine response to H1N1 and LPS in women and men.

**Conclusions and implications:** We find that STHs were associated with dampened cytokine responses to certain viral and bacterial antigens, and suggest that this mitigation of host-induced damage may reduce the incidence of cytokine storms in populations with high STH prevalence.

## 1. Introduction

Infection with soil-transmitted helminths (STHs) can modulate the host immune response and ultimately shape the outcomes (e.g., morbidity and mortality) of viral and bacterial infections [1, 2]. Because the vast majority of human history occurred in environments characterized by high STH prevalence, fundamentally shaping the evolution of human immunity, STHs may play an essential role in maintaining immunological equilibrium in the face of co-morbid infections [3, 4, 5]. Indeed, the absence of STHs in many wealthier urban communities has been linked to immune dysregulation resulting in over-active immune function and allergy [6, 5, 7], while the presence of STHs in poorer and more rural communities has been hypothesized to play a protective role against unfettered inflammatory immune responses (the ‘old friends’ hypothesis) [4, 5, 8]. Because most medical research is focused on industrialized urban environments, there are gaps in our knowledge of how STHs interact with immune function in both deleterious and potentially protective ways. STHs elicit immune responses that are polarized towards type two T helper (Th2) responses in the classical Th1/Th2 paradigm [9, 10]. Within this simplified paradigm, naive CD4+ cells are expected to differentiate into cells with distinct functions based on antigen presentation [11]. Intracellular parasites (e.g., protozoa, viruses) typically trigger Th1 responses that activate a broadly proinflammatory response (e.g., interferon gamma (IFN-*γ*), interleukin-2 (IL-2)). Helminths trigger Th2 responses that activate cells to release antiinflammatory and regulatory cytokines (e.g., IL-4, IL-5, IL-9, IL-10, IL-13, TGF-*β* ; [12, 13]. These cytokines mediate the activation of effector mechanisms that include the antibody-based immune response and regulatory T cells [14, 15, 9], inhibit the proinflammatory Th1 response, and modulate antigen reactivity [16, 17, 18]. Helminths may thus be powerful modulators of the immune response to subsequent infections. Among the most devastating outcomes of certain viral and bacterial infections is the induction of a hyperactive cytokine response (‘cytokine storm’) that can cause tissue and organ damage and is associated with many of the deaths in viral outbreaks, including the current COVID-19 pandemic [19, 8]. Cytokine storms are characterized by an unfettered increase in proinflammatory cytokines [20], and are associated with a greater incidence of severe illness and mortality [21, 19]. Because STHs can inhibit proinflammatory Th1 responses and promote regulatory T cells [22, 18], they may, paradoxically, provide protection against some of the worst outcomes of viral and bacterial infections. Understanding how STHs affect downstream viral and bacterial infections is particularly crucial for non-industrialized or rural populations with high infection prevalence. Over 12% of the world’s population is infected with at least one STH [23] and the spread of highly contagious viral and bacterial infections continues to accelerate via globalization and international travel [24]. Thus, understanding the ways in which STHs affect the immunological response to and outcomes of viral and bacterial infections is particularly exigent. This study focuses on STHs and immune response in Tsimane forager-horticulturalists of the Bolivian Amazon, who practice a largely non-industrial lifestyle centered around hunting, fishing, gathering, and small-scale slash-and-burn farming, and have relatively little interaction with market economies [25, 26, 27]. The Tsimane have high STH rates (up to 76% [28, 29]) which is likely linked to the lack of sanitation infrastructure, inconsistent access to protective footwear, shared space with animals, and open defecation practices [25, 26, 30]. We assess the hypothesis that STHs inhibit the cytokine response to viruses and bacteria by dampening the proinflammatory cytokine phenotype. We first evaluate the impact of infection with each of the five STH species most commonly found in this population on the response of 13 cytokines to whole blood sample stimulation with H1N1 vaccine and lipopolysaccharide (LPS) antigens (simulating viral and bacterial infections, respectively), predicting an association of STHs with lower proinflammatory cytokine responses. Antigens were selected based on their ability to mimic viral and bacterial infections and the feasibility of use. LPS is an abundant cell wall component of gram-negative bacteria and a frequently used antigen in stimulation studies because of its ability to induce an innate immune response characterized by inflammation [31]. The H1N1 vaccine allows for viral stimulation without using live virus in the absence of a completely controlled laboratory setting. At the time of data collection, H1N1 was not known to have spread in the area and thus would represent a novel strain that the participants had not yet encountered. We then evaluate the impact of eosinophil count, a measure of white blood cell activation that is part of the immune response to STHs, on cytokine responses to stimulation. Finally, we evaluate the impact of STHs and eosinophil count on Th1/Th2 responses, predicting that STHs and higher eosinophil counts will be associated with lower expression of a functional Th1-type category. We evaluate these patterns separately in women and men, expecting that women with STHs will exhibit higher proinflammatory responses than men based on the higher immune responsiveness (e.g., increased immune cell proliferation and activation, upregulation of immune-specific genes) baseline state observed in females across species and human populations [7, 32, 33]. All predictions are summarized in Table 1.

**Table 1.** Predictions, expected cytokine effects, and results.

| Prediction | Predicted effects | Results (H1N1 vaccine) | Results (LPS) |
| --- | --- | --- | --- |
| STH presence/absence associated with lower pro-inflammatory immune response to H1N1 vaccine and LPS stimulation | ↓ IFN-γ, IL-2<br>↓ Th1-suite<br>Women > men | Women:<br>↓ IL-1β, IL-2, IL-6, IL-7, IL-10, IL-13, GM-CSF (*A. lumbricoides* only)<br>Men:<br>none | Women:<br>↓ IFN-γ and IL-7 (*A. lumbricoides* only)<br>↓ Th1-suite<br>Men:<br>↓ IFN-γ (hookworms only) |
| Eosinophil count associated with lower pro-inflammatory immune response to H1N1 vaccine and LPS stimulation | ↓ IFN-γ, IL-2<br>↓ Th1-suite<br>Women > men | Women:<br>↓ IFN-γ, IL-1β, IL-2, IL-4, IL-7, IL-8<br>↓ Th1-suite<br>Men:<br>↓ IFN-γ, IL-2<br>↓ Th1-suite | Women:<br>↓ IFN-γ, IL-4, IL-6, IL-7, IL-8, GM-CSF<br>↓ Th1-suite<br>Men:<br>↓ IFN-γ, IL-1β, IL-6, IL-7, IL-8<br>↓ Th1-suite |

## 2. Methods

### 2.1 The Tsimane Health and Life History Project

The Tsimane Health and Life History Project (THLHP) has worked with a population of approximately 16,000 Tsimane across 90 villages since 2002, providing routine medical care and collecting epidemiological and biodemographic data [25]. Medical care is provided to all individuals and informed consent is obtained from individuals, villages, and the Tsimane government (Gran Consejo Tsimane). All research is approved by the University of New Mexico and University of California Santa Barbara Institution Research Boards (IRB # 07-157, 15-133). None of the individuals who participated in this study had received anthelmintic treatment (i.e., mebendazole, albendazole) or antibiotics from our medical staff within at least 8.5 months (mean for the 71% of sampled individuals with a record of treatment = 1.4 years) of sample collection beginning in March 2011. To our knowledge, no Tsimane had ever been vaccinated against H1N1 or other viruses at the time of this study.

### 2.2 Biomarker data collection

Biomarker data were collected from 179 Tsimane adults who participated in an antigen stimulation study. This sample included 82 women aged 29-71 (median age: 46.5) and 97 men aged 37-89 (median age: 49). Seven women were pregnant at the time of the study (based on back-calculating from time of next birth). Participants attended the THLHP Clinic in San Borja, Bolivia for routine medical care and biodemographic data collection between March and November 2011. Fasting morning blood (5 mL) was collected into a heparinized vacutainer tube. A manual white blood cell count and five-part differential (including eosinophils) were conducted with a hemocytometer immediately following each blood draw, and fecal samples were collected for parasite identification using fecal smear microscopy or a density separation technique [26].

### 2.3 Helminth infections

We characterized parasite communities by identifying parasite species with direct smear (30% of samples) microscopy [29] as well as Percoll® separation (70% of samples) in fresh fecal samples [2]. We placed all hookworm eggs in a single category because the eggs of hookworm species can be difficult to differentiate morphologically. We combined the results of both methods for all analyses, and assessed co-occurrence of parasite species with a Pearson’s correlation matrix using the ‘Hmisc’ package [34].

### 2.4 Eosinophils

Microscopic techniques such as direct smears and density separations can produce false negatives based on non-random distribution of eggs in feces and parasitic life cycles [35, 36, 37]. In addition, 30% of our fecal samples were not processed for quantitative egg counts (i.e., those that were processed with direct smears rather than Percoll separation), precluding estimation of infection intensity across the entire dataset. We thus performed additional analyses, first on the relationship between parasite infections and eosinophils, which occupy a cardinal role in the helminth-induced immune response [38] and are a common indicator of STHs in clinical settings [39, 40]. We modeled eosinophils (cells/µL) as a function of each parasite (presence/absence) for species with a prevalence of over 10% in generalized linear models (GLMs) with age included as a continuous predictor in each model (one for women, one for men). We then used eosinophil count as a proxy of parasitism in downstream analyses. As in many non-industrialized and developing populations [1], virtually no allergies or autoimmune disorders have been identified among the Tsimane [28]. The lack of allergies in such populations may arise from the regulatory immunophenotypes induced by helminth parasitism. Eosinophils are thus likely to be indicative of helminth infection in this population, although other sources of primary or secondary eosinophilia (e.g., chronic eosinophilic leukemia or certain cancers) cannot be ruled out entirely. Indeed, antihelminthic treatment is associated with significantly lowered eosinophil counts in areas with high STH infection rates [41, 42]. Eosinophils may be a useful—if coarse—metric of parasitism that captures information beyond that provided by microscopic methods. In particular, eosinophils may be most relevant during migratory larval stages characterized by pre-reproductive larvae that do not yet produce the eggs necessary to diagnose infection in fecal samples [38, 43]. Thus, we used eosinophils count here as a secondary, broader measure of parasitism that may represent, for example, infection intensity or the onset of new infections that would be less likely to be captured in fecal samples.

### 2.5 *In vitro* antigen stimulation

Aliquots of 100 µL heparinized whole blood were immediately added to separate round bottom microtiter wells in a sterile 96-well plate. One aliquot received 1µg/mL H1N1 vaccine (2009 Monovalent Vaccine (Sanofi Pasteur, Inc. Swiftwater PA 18370)) diluted in RPMI-1640. Another aliquot received 100 µL of 20 µg/mL lipopolysaccharides (LPS, Sigma cat. L2630) diluted in RPMI-1640, rendering a final concentration of 10 mg/mL LPS. Control aliquots were run in RPMI alone. To prevent contamination, RPMI was supplemented with 100 IU/mL penicillin and 100 µg/mL streptomycin (Sigma cat. P0781), and any plates with visible growth or excessive RPMI values that might indicate non-visible contamination were eliminated. We enriched CO_2_ concentration by sealing plates in an airtight container with a burning candle to deplete O_2_ as a field-friendly alternative to a CO_2_ incubator [44]. The sealed and treated blood samples were incubated at 37°C for 72 hours. Samples were centrifuged and supernatants were frozen in liquid nitrogen, transported on dry ice, and stored at -80°C for up to 2 years (see [45, 46]). At the Hominoid Reproductive Ecology Laboratory at the University of New Mexico, 13 cytokines (INF-*γ*, IL-1*β*, IL-2, IL-4, IL-5, IL-6, IL-7, IL-8, IL-10, IL-12p70, IL-13, Granulocyte-macrophage colony-stimulating factor (GM-CSF), and Tumor necrosis factor-alpha (TNF-*α*)) were measured with a Milliplex MAP High-Sensitivity Human Cytokine Panel (HSCYTMAG-60SK-13, Millipore Corp., Billerica, MA) on a Luminex MagPix (Millipore Corp., Billerica, MA). All quality control specimens were within normal limits.

### 2.6 Statistical analysis

#### 2.6.1 Effect on individual cytokine responses

We first assessed the impact of STHs on the response of each cytokine to stimulation with H1N1 and LPS antigens. Modeling women and men separately, we used GLMs to model the log-transformed concentration of each cytokine in either H1N1 of LPS as a function of the presence or absence of each STH. We included body mass index (BMI) and age as covariates in all models, and, in the female specific models, included a covariate for pregnancy (0/1) [47]. While previous research in the Tsimane has shown differences in immune function by trimester, the small number of pregnant women (n=6) in this study precluded a trimester-level analysis [48]. We included in our models only STHs with a prevalence of over 10% across our sample set. We then assessed the impact of eosinophil count (cells/µL) on individual cytokine response to stimulation. We again used GLMs to model the response of log-transformed cytokines in response to stimulation with H1N1 and LPS as a function of eosinophil count. We included BMI, age, and pregnancy for women, and included total leukocyte count to account for variation in total white blood cells. We assessed collinearity between eosinophil and leukocyte counts by examining the variation inflation factor (VIF) for all models; all VIFs were acceptably low (<2.5). We found no significant interaction between age and eosinophils and thus omitted this interaction from our final models. We calculated a meta effect score by combining the fixed effects from each model for women and men (‘meta’ package; [49]). Although this approach assumes cytokine independence, it allows for a broad estimation of the cross-cytokine effect of eosinophils and functions as a summary statistic. We did not consider changes from baseline cytokine levels (unpublished data) because cytokines have short half-lives, ranging from minutes to hours [50]; thus, many of the circulating cytokines would have degraded during incubation and would not be present at biological relevant levels at the time of assay.

#### 2.6.2 Effect on Th1-Th2 response

Because cytokines are pleiotropic, we created functional Th1 and Th2 categories according to standard designations. Generally, Th1-type cytokine groups include proinflammatory cytokines such as IFN-*γ* and IL-2 [51], but researchers have also included IL-6, IL-12p70 and TNF-*α* in this category [52]. Likewise, Th2-type cytokine groups generally include anti-inflammatory or regulatory cytokines such as IL-4, IL-5, and IL-13 [51], though researchers have also included IL-10 [52]. Our categories were based on the most conservative designations because many cytokines can play multiple roles in different contexts; our Th1-suite included IFN-*γ* and IL-2, while the Th2-suite included IL-4, IL-5, and IL-13 [51]. We ran parallel analyses with broader categories (included in the supplementary materials; Table S10). In the proinflammatory suite, we included IFNy, IL-2, IL-6, IL-8, IL-12p70, and TNFa; in the anti-inflammatory suite, we included IL-4, IL-5, IL-10, IL-13. We added a constant (+1) to each cytokine value, which we then log-transformed. We calculated z-scores for each of the functional suites within each treatment with the sum of the individual cytokines. Finally, we averaged z-scores within each category to produce z-scored suite responses for each sample. Assessing women and men separately, we first modeled z-scores within each treatment as a function of each STH (presence/absence), BMI, age, and pregnancy (for women). We then performed similar analyses modeling the effect of eosinophil count on Th1/Th2-suite z-scores, including the same covariates as predictors as well as total leukocyte count. All analyses were done in R.

## 3. Results

### 3.1 Descriptives

Helminth richness ranged from 0-4 species per person (median: 1, Figure 1, n=179), and 87% of samples had at least one helminth infection. We identified hookworm (*Necator americanus* or *Ancylostoma duodenale*) (75.4% prevalence), *Ascaris lumbricoides* (21.8%), *Strongyloides stercoralis* (13.4%), *Trichuris trichiura* (5.6%), and *Hymenopolis nana* (0.6%). The protozoan *Giardia lamblia* (13.4%) and the amoeba *Entamoeba histolytica* (7.3%) were also identified. Coinfections (37%) were most likely to occur with *A. lumbricoides* and *T. trichiura* (Pearson correlation = 0.23, p < 0.01, Table S1) and least likely to occur between *A. lumbricoides* and hookworms (Pearson correlation = -0.27, p < 0.01, Table S1). Only helminths with a prevalence of >10% were included for downstream analyses (*A. lumbricoides, S. stercoralis*, and hookworm species; Table S2).

**Figure 1.**
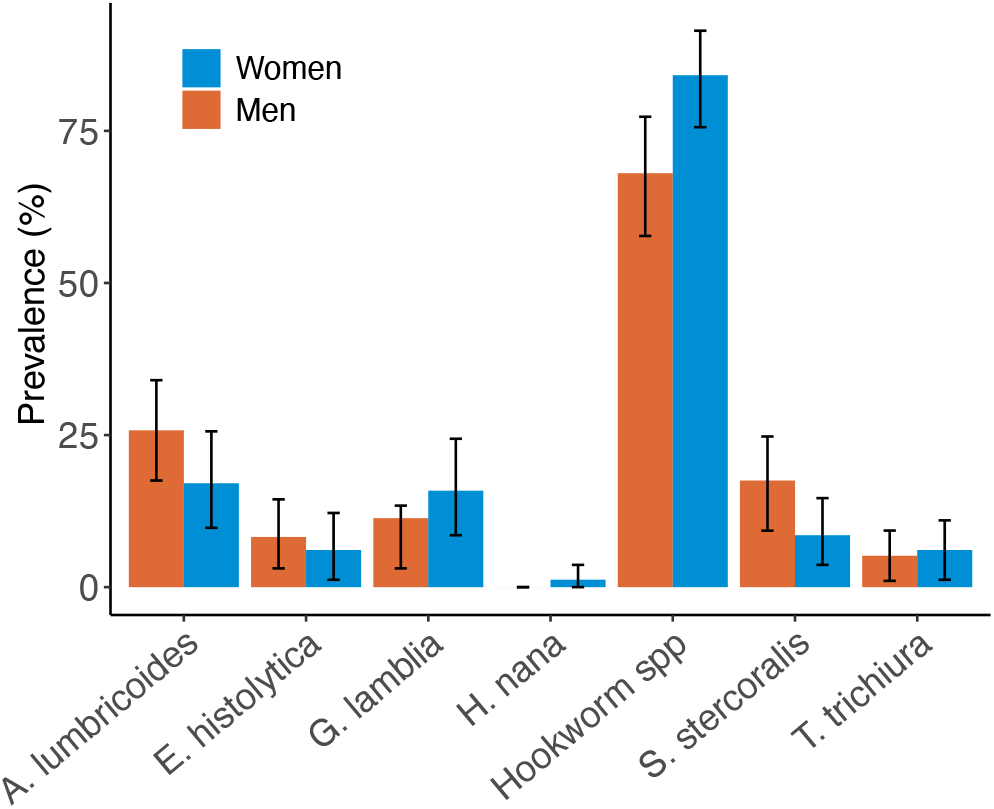
Prevalence of parasites identified microscopically in 179 fresh fecal samples.

Eosinophil counts ranged from 492 - 7,200 cells/µL, with a median of 1,950 cells/µL and a mean of 2,230 cells/µL. For comparison, the US reference range for eosinophils is <500 cells/µL; >500 cells/µL is categorized as eosinophilia [53]. Thus, by US standards all but two (98.6%) of the sampled Tsimane were eosinophilic.

### 3.2 Parasites and eosinophils

The presence of *S. stercoralis* was the only parasite infection positively associated with increased eosinophils (p = 0.004) for men. No parasite infection was associated with changes in eosinophil count for women.

### 3.3 Individual cytokine responses

#### 3.3.1 H1N1 stimulation

For women, the presence of *A. lumbricoides* was significantly (p ≤0.05) negatively associated with the response of seven cytokines to H1N1 (IL-1*β*, IL-2, IL-6, IL-7, IL-10, IL-13, and GM-CSF; Table S4). *A. lumbricoides* was the sole parasite to exhibit a significant relationship with cytokine responses for women, and exhibited negative betas associated with all 13 cytokines (Figure 2, Table S4). The overall estimate of the impact of *A. lumbricoides* across cytokines for women in the fixed-effects model was -0.6 (p < 0.01; Figure 2, Table S4). Eosinophils were significantly negatively associated with the response of six cytokines to H1N1 (IFN-*γ*, IL-1*β*, IL-2, IL-4, IL-7, IL-8), with eosinophils exhibiting negative betas for all of the 13 total cytokines but IL-5 (Figure 2, Table S5). The overall estimate of eosinophil impact across cytokines for women in the fixed-effects model was -0.8 (p < 0.01; Figure 2, Table S4). Age was positively associated with TNF-*α* and GM-CSF responses to H1N1 in parasite-specific models (Table S5). For men, individual parasite infections were not significantly associated with cytokine responses to H1N1 stimulation (Table S6). Eosinophils were significantly negatively associated with the response of two cytokines to H1N1 (IFN-*γ* and IL-2), and exhibited negative betas for all cytokines but IL-5 (Figure 2, Table S7). The overall estimate of eosinophil impact across cytokines in the fixed-effects model was -0.63 (p < 0.01; Figure 2, Table S7). Age, BMI, pregnancy, and leukocyte count had varying effects on cytokine response to H1N1 in all models (Figure 2, Table S7).

**Figure 2.**
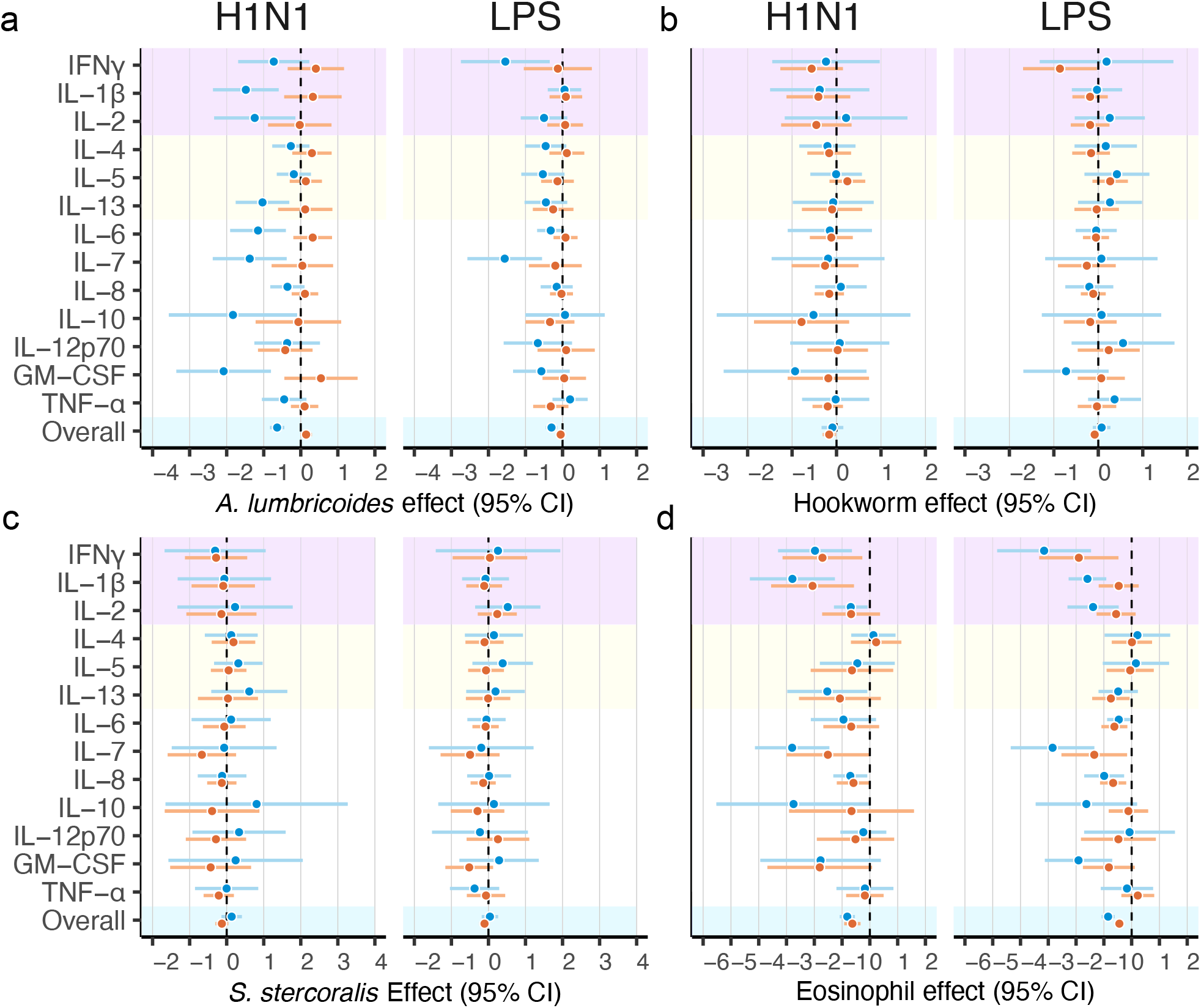
Individual cytokine responses to H1N1 and LPS stimulation as a function of (a) *A. lumbricoides*, (b) hookworms, (c) *S. stercoralis*, and (d) eosinophil count. Coefficients and 95% confidence intervals are shown for each condition for women (blue) and men (orange). Cytokines are shaded by their Th1- (purple) or Th2-type (yellow) classification, and the overall effect size for each stimulation is highlighted in blue.

#### 3.3.2 LPS stimulation

For women, the presence of *A. lumbricoides* was significantly negatively associated with the response of two cytokines to LPS stimulation (IFN-*γ* and IL-7; Table S4). No other parasite infection demonstrated any relationship with cytokine expression. The overall estimate of the impact of *A. lumbricoides* on cytokine response in the fixed-effects model was -0.3 (p < 0.01; Table S4). Eosinophils were significantly negatively associated with the response of seven cytokines to LPS (IFN-*γ*, IL-4, IL-6, IL-7, IL-8, and GM-CSF), and produced negative betas for all cytokines except IL-5 and IL-13 (Figure 2, Table S5). The overall estimate of eosinophil impact across cytokines in the fixed-effects model was -0.9 (p < 0.01; Table S5). Age, BMI, pregnancy, and leukocyte count had varying effects on cytokine responses to LPS. Age was positively associated with TNF-*α* and GM-CSF responses to LPS in parasite-specific models (Table S5). Additional models excluding pregnant women were not substantially different from the primary models. For men, the presence of hookworms was significantly negatively associated with the expression of IFN-*γ* to LPS stimulation (Table S6). No other parasite infection exhibited any relationship with cytokine expression. Eosinophils were significantly negatively associated with the response of five cytokines to LPS (IFN-*γ*, IL-1*β*, IL-6, IL-7, and IL-8), and produced negative betas for all cytokines except IL-5 and TNF-*α* (Figure 1, Table S7). The overall estimate of eosinophil impact across cytokines in the fixed-effects model was -0.44 (p < 0.01) for men (Figure 1), and age, BMI, and leukocyte count had varying effects on cytokine response to LPS. Specifically, age was associated with higher IL-7 response to LPS stimulation (Table S7).

### 3.4 Th1-Th2 responses

For women, *A. lumbricoides* was not associated with the Th1- or Th2-type cytokine suite to H1N1 stimulation, and was negatively associated with the response of the Th1-type cytokine suite (*β* = 0.6, p = 0.04) to LPS stimulation but not with the response of the Th2-type cytokine suite (Table S8). Eosinophil counts were negatively associated with the response of Th1-type (*β* = -1.06, p < 0.01) but not Th2-type (*β* = -0.43, p = 0.13) cytokines to H1N1, with the same pattern observed in response to LPS stimulation (Th1-type: *β* = -1.45, p < 0.01, Th2-type: *β* = -0.41, p = 0.47, Figure 2, Table S9). For men, no parasites were significantly associated with the response of either cytokine suite to stimulation with either H1N1 or LPS. Eosinophil counts were negatively associated with response of Th1-type (*β* = -1.13, p = 0.01) but not Th2-type cytokines (*β* = -0.3, p = 0.54; Figure 2, Table S8) to H1N1 stimulation. Similarly, eosinophil counts were associated with the response of Th1-type (*β* = -0.64, p = 0.05) but not Th2-type cytokines (*β* = -0.22, p = 0.51; Figure 2, Table S9) to LPS stimulation. The other predictors had varying effects on the expression of Th1- and Th2-type cytokines for women and men. Higher age was associated with higher Th1 responses to H1N1 for women (a similar effect was observed for LPS but it did not meet the significance threshold); no association was observed in men in either media (Table S9).

## 4. Discussion

As suggested by evolutionary hypotheses (e.g., “old friends”), STH infections (specifically, *A. lumbricoides*) are associated with dampened proinflammatory responses to acute viral and bacterial stimulations among Tsimane women. Eosinophils, considered here as a secondary indicator of infection status, were also significantly associated with lower cytokine responses to H1N1 and LPS in women and men. Together, these results suggest that helminth infections may attenuate the proinflammatory response to viruses and bacteria with the strongest effect in women.

### 4.1 Helminths may dampen the acute immune response to infection

STH infection in the Tsimane was high, with at least one helminth found in 87% and two or more helminths found in 37% of sampled individuals. In conjunction with the high levels of immunoglobulin E (IgE) observed in the Tsimane [26, 28], the high eosinophil levels suggest that the prevalence of STHs is likely even higher than those identified microscopically. The most common helminth infections were with the gastrointestinal nematodes *A. lumbricoides, S. stercoralis, T. trichiura*, and hookworms, all of which generally elicit anti-inflammatory and regulatory immune responses characterized primarily by Th2 cytokine cascades [54, 55, 56], with some evidence for a role of Th1 cytokine responses in certain of these infections [55]. These primarily anti-inflammatory immune responses can inhibit the ability of the immune system to launch pro-inflammatory responses in the face of viral and bacterial pathogens. The pro-inflammatory immune responses elicited by H1N1 and similar viruses are characterized by elevations in IFN-*γ*, IL-1, IL-2, IL-5, IL-6, IL-8, IL-12, and TNF-*α*, among others [57]. While a certain degree of responsiveness is vital to combating infection, a proinflammatory ‘cytokine storm’ can culminate in tissue and organ damage; indeed, deaths attributed to viral infections and certain gram-negative bacteria (e.g., E. coli [21, 58]) are typically associated with damages arising from cytokine storms [20, 19]. In Tsimane women, infection with *A. lumbricoides* was significantly associated with impaired responses of certain cytokines implicated in cytokine storms [59, 57]: namely, IFN-*γ* (in LPS stimulation) and IL-1*β*, IL-2, and IL-6 (in H1N1 stimulation). This suggests that Tsimane women infected with *A. lumbricoides* may be less susceptible to the development of cytokine storms during the trajectory of viral or bacterial infections. In Tsimane men, infection with hookworms was only associated with a diminished response of IFN-*γ* in LPS. In the Th1/Th2-suite analyses, the only observed effect was the dampening of the Th1-suite response to LPS in women by *A. lumbricoides*. The sex-specific effect of *A. lumbricoides* occurs despite fairly equal infection prevalence among women and men (Table 1), and may be tied to parasite-specific sex differences in immunity [60, 61] and the profiles of chronic versus acute infections. As expected based on both the limitations of traditional microscopy and the binary nature of our parasite infection data, eosinophils were negatively associated with more cytokines than were individual parasite species. In women, higher eosinophil count was associated with decreased expression of both pro- and anti-inflammatory cytokines: namely, IL-4, IL-6, IL-7, IL-8 and GM-CSF (in LPS stimulation) and IL-1*β*, IL-2, IL-6, IL-7, IL-10, and IL-13 (in H1N1 stimulation). In men, the effect was similarly spread across anti- and pro-inflammatory cytokines. In the functional Th1/Th2-suite analyses, eosinophils were associated with inhibited expression of the Th1-suite for both women and men in all media. Associations between eosinophils and cytokine expression diverged from those between parasite presence and cytokine expression (i.e., certain cytokines were associated with eosinophils, but not with a specific parasite, and vice versa). This may be due to the limitations of our parasite detection techniques, or may be tied to factors such as infection intensity that were not explicitly quantified here but may be reflected in our eosinophil measure. In addition, eosinophils may be most relevant for infections at their onset for the specific suite of parasites included in these analyses. Infections with *A. lumbricoides, S. stercoralis*, and hookworms all involve an initial larval migration; our analyses thus may capture infections at various points along the infection timeline. Eosinophil count may thus reflect infections at their onset, prior to the reproductive phase characterized by high egg counts that would be most likely to be captured with microscopy. Eosinophils may also protect against new infections, represent the tail-end of recently cleared infections, or reflect early life exposure to helminths that would prime individuals for anti-inflammatory responses throughout their lives. These components of STH parasitism may underlie the relationship observed between antihelminthic treatment and eosinophil count in certain populations with high STH infection rates [42, 41]. Indeed, Tsimane have significantly elevated IgE by the age of 5, and cross-sectional studies suggest that IgE remains high across the lifecourse [2] consistent with lifetime STH exposure and infection. These results suggest that eosinophils may reflect a to-be-determined component of STH infection and recapitulate the immunosuppressive associations in the STH analyses. We found the strongest immunosuppressive associations in women, pointing to sex differences in immune function deviating from the expected pattern of immune hyperactivity in women [7]. Women are generally less susceptible to pathogens [62, 63] and mammalian females tend to host parasite infections at lower rates and with lower loads than males [64] (but see [65]), but have historically incurred higher morbidity and mortality associated with viral pandemics (reviewed in [66]). These patterns have been linked to the proinflammatory properties of estradiol [66] and the immunosuppressive components and energetic costs of testosterone [67, 45]. Much of the data that have pointed to higher morbidity and mortality in women during viral pandemics, however, are from industrialized populations, which are typically characterized by low STH rates [3], as well as high autoimmune risk, and low fertility [7]. Our results suggest that concurrent STH infections may mitigate the risk of overreaction to viral and bacterial infection and may buffer women from the otherwise higher likelihood of development of cytokine storms through suppression of cytokine responses.

**Figure 3.**
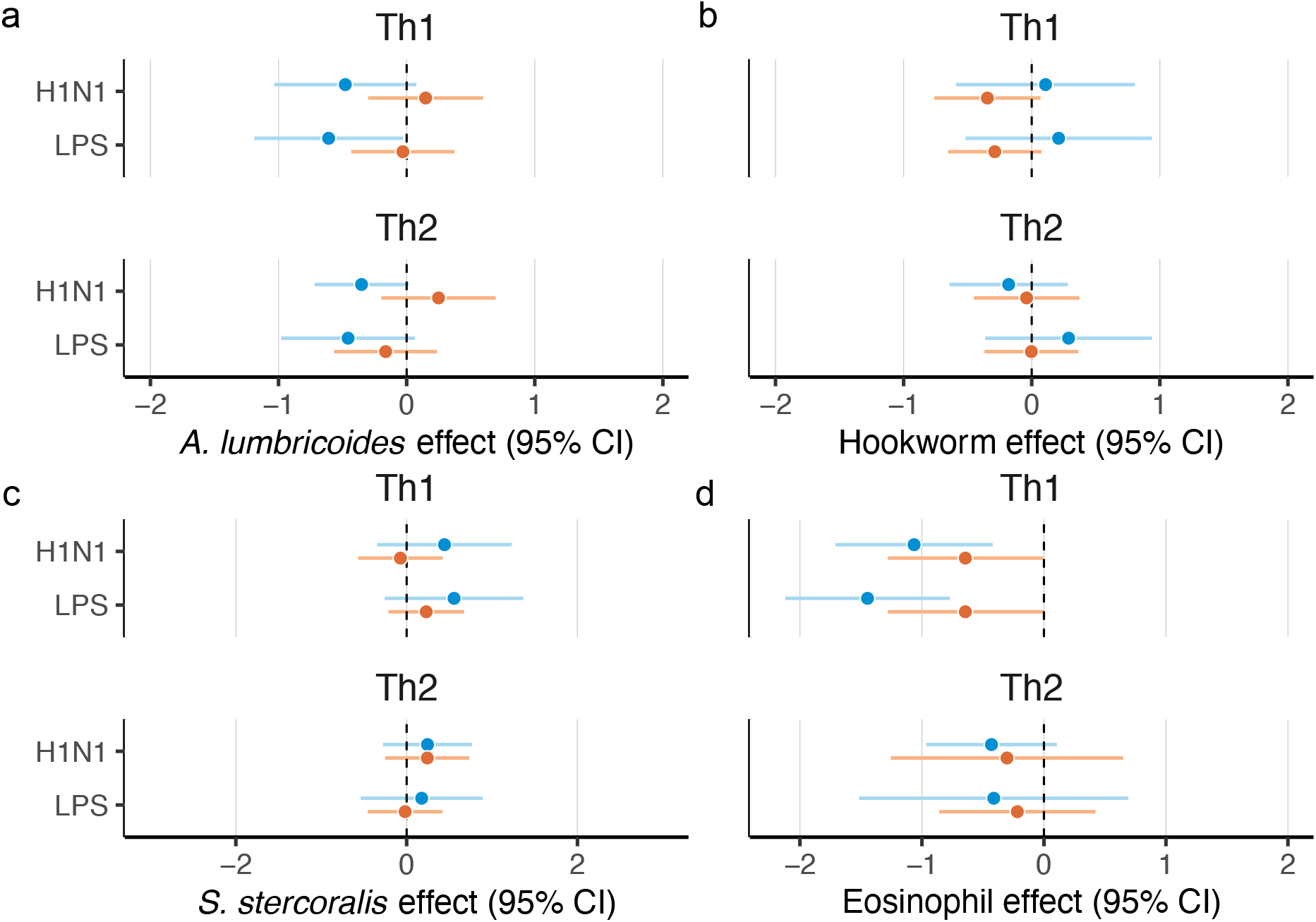
The effect of (a) *A. lumbricoides*, (b) hookworms, (c) *S. stercoralis*, and (d) eosinophils on Th1- and Th2-type responses to LPS and H1N1 stimulation. Coefficients and 95% confidence intervals (CIs) are shown for each condition for women and men.

### 4.2 Implications for COVID-19

Beyond attenuating the overactive immune responses that underlie allergies, heart disease, and diabetes [5, 68, 1], the helminth-induced anti-inflammatory immunomodulatory network may also attenuate some of the most severe symptoms of SARS-CoV-2 infection in populations with high helminth infection prevalence [8, 4]. While we did not assess the impact of SARS-CoV-2 stimulation on cytokine responses, our viral stimulation suggests that STHs may be protective against septic shock and mortality characteristic of COVID-19. Heterogeneity in hyperactive immune responses to viruses is a major predictor of health outcomes, and some of this may derive in part from environmental exposure to STHs. This would suggest that morbidity and mortality due to COVID-19 may be reduced in areas with high helminth prevalence, which has been discussed as a contributor to early heterogeneity in global COVID-19 mortality [69]. Other components of health that may contribute to this pattern are cardiovascular disease and obesity, both of which have been linked to COVID-19 mortality [70]. Notably, the Tsimane have the lowest rates of cardiovascular disease reported for any population [71]. Global COVID-19 mortality patterns are likely the product of complex webs of community-specific demographic, anthropometric, nutritional, and behavioral factors, interacting with individual-level immunocompetence, that will continue to be unraveled as the pandemic progresses.

### 4.3 Limitations

Because of the lack of standard health surveillance in the region, we are unable to know for certain if study participants had previously been exposed to H1N1, which would shape the interpretation of our results. We know that immunological changes during pregnancy vary by trimester [48], but our low sample size of pregnant women precluded a trimester-level analysis. We also did not include children in our analyses and thus cannot assess the role of early life exposure in this relationship. Finally, this study is correlational and did not assess the causality underlying the observed relationships between STH infections, eosinophils, and immune function. Nevertheless, studies that include data from non-industrialized populations are relatively rare and provide knowledge that would not be possible to produce in a laboratory.

### 4.4 Conclusions

Altogether, our results support the hypothesis that STHs inhibit the cytokine response to viruses and bacteria by specifically dampening the proinflammatory cytokine phenotype. These findings are consistent with those that have observed lower immunogenicity of vaccines in people with helminth infections [72] and support the dampening effect of helminths on the proinflammatory response. However, we also observed associations between STHs and cytokines that are generally characterized as anti-inflammatory or can express anti-inflammatory properties (IL-4, IL-6, IL-10, IL-13). It is thus possible that the effect of STHs on immunity stretches beyond effects on the proinflammatory immune responses, and may work through indirect mechanisms such as STH-induced nutritional deficiencies or anemia. Notably, our results also suggest that sex-based differences in immune function described in other populations and species [33, 64] hold true in the Tsimane, with women exhibiting stronger immune responses to stimulations. Further research can increase the granularity of this knowledge by using cell sorting to determine which cytokine-producing cell types (e.g., B-cells, T-cells, macrophages, leukocytes) are affected. Overall, these results add to a growing level of support for the “old friends” hypothesis, suggesting that STH infections may play an essential role in maintaining immunological equilibrium in the face of downstream infections [3, 4, 5].

## Acknowledgements

We thank the Tsimane participants and communities who engaged in this research and the Tsimane Health and Life History Project staff and personnel for their efforts and logistical support. We thank Thais Mendes for laboratory assistance, and Maria Yazdanbakhsh, E rliyani Sartono, and Linda May for assistance with stimulation protocols, and we thank Kenny Chiou for visualization assistance. This research was funded by NIH/NIA (R01AG024119, RF1AG054442) and the ASU Center for Evolution and Medicine. JS Stieglitz acknowledges IAST funding from the French National Research Agency (ANR) ANR-17-EURE-0010 (Investissements d’Avenir program).

